# Niche-derived soluble DLK1 promotes glioma stemness and growth

**DOI:** 10.1101/2020.08.20.258608

**Authors:** Elisa S. Grassi, Pauline Jeannot, Vasiliki Pantazopoulou, Tracy J. Berg, Alexander Pietras

**Affiliations:** Division of Translational Cancer Research, Department of Laboratory Medicine, Lund University, Lund, Sweden

**Keywords:** DLK1, hypoxia, HIF-2alpha, glioma, tumor-associated astrocytes, stem cell niche, tumor progression

## Abstract

Tumor cell behaviors associated with aggressive tumor growth such as proliferation, therapeutic resistance, and stemness are regulated in part by soluble factors derived from the tumor microenvironment. Tumor-associated astrocytes represent a major component of the glioma tumor microenvironment, and astrocytes have an active role in maintenance of normal neural stem cells in the stem cell niche, in part via secretion of soluble Delta-like Non-Canonical Notch Ligand 1 (DLK1). We found that astrocytes, when exposed to stresses of the tumor microenvironment such as hypoxia or ionizing radiation (IR), increased secretion of soluble DLK1. Tumor-associated astrocytes in a glioma mouse model expressed DLK1 in perinecrotic (hypoxic) and perivascular tumor areas. Glioma cells exposed to recombinant DLK1 displayed increased proliferation, enhanced sphere and colony formation abilities, and increased levels of stem cell marker genes. Mechanistically, DLK1-mediated effects on glioma cells involved increased and prolonged stabilization of Hypoxia-Inducible Factor 2alpha (HIF-2alpha), and inhibition of HIF-2alpha activity abolished effects of DLK1 in hypoxia. Forced expression of soluble DLK1 resulted in more aggressive tumor growth and shortened survival in a genetically engineered mouse model of glioma. Together, our data support DLK1 as a soluble mediator of glioma aggressiveness derived from the tumor microenvironment.

**Graphical abstract:** 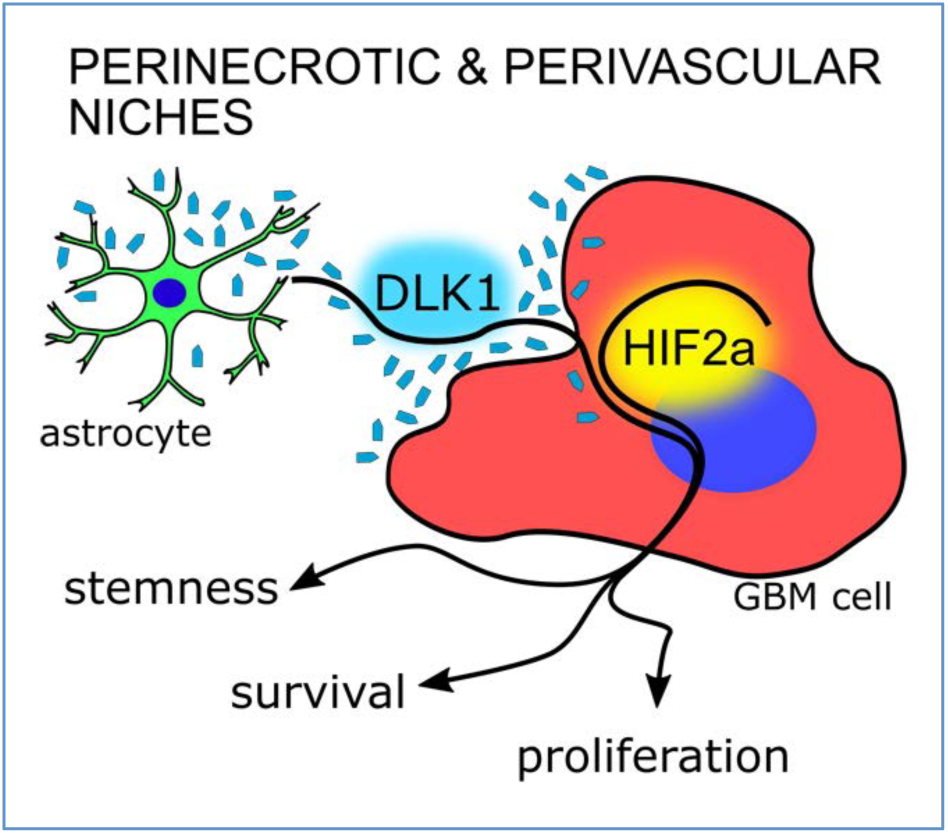

**Highlights:**

- Astrocytes secrete DLK1 after exposure to hypoxia or irradiation
- Soluble DLK1 promotes stemness in glioma, in part by increasing HIF-2alpha stabilization.
- High levels of soluble DLK1 are associated with tumor aggressiveness and lethality.

## INTRODUCTION

Glioblastoma recurrence following standard-of-care treatment with radiotherapy, surgery, and chemotherapy invariably gives rise to incurable lesions, and the median survival following diagnosis remains dismal at 15-20 months despite recent advances in our understanding of glioblastoma at a molecular level (Huse & Holland, 2010). Evidence from human patient samples and murine models of brain tumors suggest that inherent therapeutic resistance within a subset of tumor cells with stem cell characteristics may be the primary source of recurrent tumors (Bernstock et al., 2019; Dirkse et al., 2019). While the origin and fate of such cells remain controversial, glioblastoma cell phenotypes are highly plastic, and non-stem-like cells can acquire characteristics of stem cells as a result of microenvironmental interactions with the extracellular matrix, with growth factors, or as a result of altered oxygenation or pH (Dirkse et al., 2019; Hambardzumyan & Bergers, 2015; Jawhari, Ratinaud, & Verdier, 2016; Ryskalin et al., 2019). Indeed, tumor cells with stem cell properties appear to be spatially restricted to specific microenvironments such as the perinecrotic (hypoxic) and perivascular niches, suggesting that these niches may control residing tumor cell phenotypes (Hambardzumyan & Bergers, 2015; Jawhari et al., 2016; Majmundar, Wong, & Simon, 2010).

Delta Like Non-Canonical Notch Ligand 1 (DLK1) is a transmembrane protein in the Notch family of ligands, that is capable of signaling in a Notch-dependent and – independent manner depending on cellular context (Baladrón et al., 2005; Ceder, Jansson, Helczynski, & Abrahamsson, 2008; Falix, Aronson, Lamers, & Gaemers, 2012; Ferrón et al., 2011; Huang et al., 2019; Li et al., 2014). Expression of DLK1 is increased with tumor grade in glioma, and its signaling has been associated with various tumor cell properties such as proliferation, invasion, and stemness (Grassi, Pantazopoulou, & Pietras, 2020; Yin et al., 2006). The mechanisms underlying these effects on tumor cell behavior remain poorly understood, but likely include signaling from the extracellular, soluble domain of DLK1 (Wang & Sul, 2006). Indeed, soluble DLK1 secreted from astrocytes was recently shown to be a critical component of the subventricular zone (SVZ) neural stem cell niche (Ferrón et al., 2011). Astrocytes represent a prominent cell type in the brain tumor microenvironment (Mega et al., 2020), and recent studies revealed that DLK1 is one of the top upregulated genes in tumor-associated astrocytes of high-grade vs. low-grade gliomas (Katz et al., 2012). The regulation of DLK1 expression is poorly understood, but some elements of the brain tumor microenvironment such as hypoxia have been shown to drive DLK1 expression (Begum, Kim, Lin, & Yun, 2012; Kim, Lin, Zelterman, & Yun, 2009).

Here, we sought to investigate the role of soluble DLK1 in the high-grade glioma tumor microenvironment. We found increased secretion of DLK1 from tumor-associated astrocytes subjected to stresses of the tumor microenvironment, such as hypoxia and ionizing radiation (IR). Soluble DLK1 increased proliferation and stemness of glioma cells, and promoted tumor growth in a genetically engineered mouse model of glioma. Together, our findings suggest that soluble DLK1 is a niche-derived mediator of aggressive tumor growth in brain tumors.

## MATERIALS AND METHODS

### Glioma mouse models

Nestin-tv-a (Ntv-a) Ink4a/Arf^−/−^ mice (IMSR Cat# NCIMR:01XH4, RRID:IMSR_NCIMR:01XH4) were intracranially injected with DF1 cells (ATCC Cat# CRL-12203, RRID:CVCL_0570) expressing RCAS-PDGFB and RCAS-shp53, as indicated, in the neonatal brain with a 10-μL gas-tight Hamilton syringe, as described previously (Bleau et al., 2009; Holland, Hively, DePinho, & Varmus, 1998).

Soluble DLK-S was cloned into the RCAS vector and mice were co-injected with a 1:1 mix of DF1 cells expressing PDGFB and DLK-S or empty RCAS as indicated. Each litter was allocated to one experimental group. Mice were monitored daily and sacrificed upon displaying brain tumor symptoms. All procedures were approved by the Malmö-Lund Ethical Committee (permit M186-14). The sample number was determined based on the law of diminishing returns with the resource equation method (total number of animals – total number of groups >10). A total of 6 pups were excluded due to non-tumor symptoms during week 0-3, the final numbers were n=26 for PDGFB and n=24 for DLK-S.

### Immunofluorescence

Whole brains were embedded in OCT (ThermoFisher) and frozen in pre-cooled isopentane. 5 µm thick cryosections were air-dried for 30 min, then fixed in ice-cold acetone or 4% PFA and permeabilized using 0.3% Triton X-100 in PBS (Sigma). Blocking was performed using serum-free protein block (DAKO), then sections were incubated overnight with primary antibodies at 4°C with Background Reducing Components (DAKO). Alexa Fluor secondary antibodies (Abcam) were used, and Vectashield Mounting medium with DAPI (Vector Laboratories) was used for mounting.

Primary antibodies: Pref-1/DLK1/FA1 Antibody (Novus, Cat# NBP2-33697), DLK1 Polyclonal Antibody (Thermo Fisher Scientific Cat# PA5-72199, RRID:AB_2718053), Mouse Pref-1/DLK1/FA1 Antibody (R and D Systems, Cat# AF8277), HIF2 alpha antibody (Abcam Cat# ab199, RRID:AB_302739), Goat Anti-Human Olig2 (R and D Systems Cat# AF2418, RRID:AB_2157554), Chicken Anti-GFAP (Abcam Cat# ab4674, RRID:AB_304558), Ki-67 antibody (Thermo Fisher Scientific Cat# RM-9106-S0, RRID:AB_2341197).

Secondary antibodies: Donkey Anti-Goat IgG H&L Alexa Fluor 555 (Abcam Cat# ab150134, RRID:AB_2715537), Donkey anti-Rabbit IgG (H+L) Alexa Fluor 488 (Thermo Fisher Scientific Cat# A-21206, RRID:AB_2535792), Donkey anti-Chicken IgY H&L FITC (Abcam Cat# ab63507, RRID:AB_1139472) Goat anti-Rabbit IgG (H+L) Alexa Fluor 568 (Thermo Fisher Scientific Cat# A-11011, RRID:AB_143157).

Images were acquired using an Olympus BX63 microscope and DP80 camera and CellSens software (Olympus CellSens Software, RRID:SCR_016238).

For DLK1 and GFAP localization images (Fig. 1), minimal postproduction consisting of background subtraction and automated level optimization was equally applied with ImageJ (Fiji, RRID:SCR_002285).

**Figure 1.**
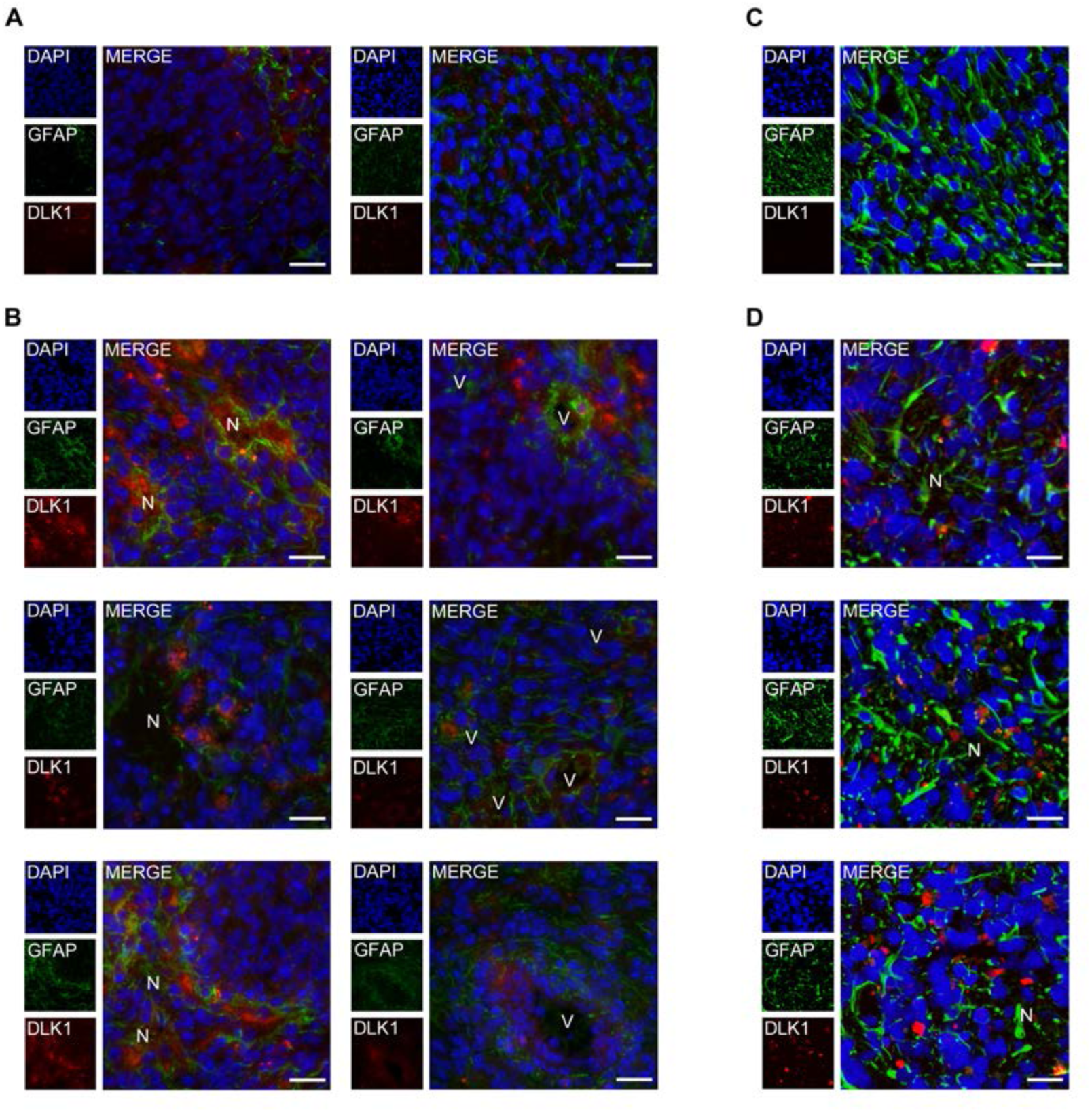
A, B: Representative images of immunofluorescent stainings showing tumor-associated astrocytes and DLK1 localization in bulk tumor (A) and perinecrotic (N) and perivascular (V) areas (B) of shp53-induced murine gliomas. Scalebars represents 25µm. C,D: Representative images of immunofluorescent stainings showing tumor-associated astrocytes and N-terminal, secreted, DLK1 localization in bulk tumor (C) and perinecrotic areas (N)(D) of shp53-induced murine gliomas. Scalebars represents 25µm. Statistical analysis: A, B, C, D n=3

Colocalization analysis was performed with ImageJ (Fiji, RRID:SCR_002285) Coloc2 plugin on the selected Regions of Interest of at least 3 independent experiments.

Ki67 quantification was performed with CellProfiler (CellProfiler Image Analysis Software, RRID:SCR_007358), at least 3 fields were analyzed for each tumor, for a total of 101717 nuclei analyzed, with n=45466 for PDGFB tumors and n=56251 for DLK-S tumors.

### Cell culture and treatments

Primary murine glioma cells (PIGPCs (Johansson et al., 2017)) were cultured as described in DMEM (Life Technologies) + 10% fetal bovine serum (FBS) and 1% PenStrep (Corning). U3082MG (RRID:CVCL_IR93), U3084MG (RRID:CVCL_IR94) and U3065MG (RRID:CVCL_IR87) cells were from A Human Glioblastoma Cell Culture Resource (HGCC) and were cultured as described (Xie et al., 2015) in Neurobasal medium (GIBCO)/DMEM/F12 with Glutamax (Life Technologies) +1% PenStrep solution, N2 and B27 (Life Technologies), 10 ng/mL epidermal growth factor (EGF), and 10 ng/mL fibroblast growth factor (FGF) (Peprotech). Cells were dissociated using Accutase (ThermoFisher), and cultured as monolayers in dishes coated with polyornithine (Sigma) and laminin (Biolamina). Primary Human Astrocytes (3H Biomedical Cat# 1800-10) were cultured in Human Astrocyte Medium #SC1801 (3H Biomedical) and were used below passage 15. A Gamma Cell-3000 Elan (MDS Nordion) was used to deliver 10 Gy in a single dose. A Whitney H35 Hypoxystation (Don Whitley Scientific) was used to generate hypoxic conditions.

Cells were treated with the indicated concentrations of DLK1 human (Sigma Cat# SRP8006) or Recombinant Human DLK-1 protein (Abcam Cat# ab151926). For HIF2alpha inhibition, cells were treated with 10µM PT2385 (MedChemExpress Cat# HY-12867) or equivalent amount of DMSO 24h prior hypoxia exposure and kept in the dark for the whole experiment. Transient transfection was performed with X-tremeGENE 9 DNA Transfection Reagent (Sigma Cat# 06 365 809 001) following manufacturer’s indications. Human DLK1 ectodomain expression vector was obtained from Addgene (DLK1-bio-His, RRID:Addgene_51876) (Sun et al., 2015).

### DLK1 ELISA assay

DLK1 secretion was measured with Human Pref-1 ELISA kit for cell culture supernatants, plasma and serum samples RAB 1076 (Sigma-Aldrich). Medium from control and treated astrocytes was collected every 2-3 days while medium from DLK-S transfected cells was collected 72h hours post transfection, immediately snap frozen in dry ice and stored at −80°C until the day of the assay. The ELISA was performed following manufacturer’s instructions and 450 nm absorbance was read on a Synergy 2 plate reader (BioTek).

### Proliferation assay

1000 cells/well (PIGPC) or 2500 cells/well (U3082MG, U3084MG and U3065MG) were seeded in 96 well plates. For astrocyte conditioned media (ACM) transfer experiments, 24 hours after seeding, media from Astrocyte and transfected cells was filter sterilized and used to replace culture media. For astrocyte experiments, media was replaced every 2-3 days and proliferation was assessed after 9 days. For transfected cells, proliferation was assessed after 72 hours.

For recombinant protein experiments, 24 hours after seeding cells were treated with serial dilutions of recombinant DLK1 (0-200 ng/ml range) and grown for 72 hours. At the moment of the assay, 10 µl of WST-1 solution (Roche) were added to each well and after 2 hours of incubation at 37°C and 5% CO2 450 nm absorbance was read on a Synergy 2 plate reader (BioTek).

### Western Blot

Cells were lysed in RIPA buffer supplemented with Complete Phosphatase and Complete Protease inhibitor cocktails (Roche). After dilution in Laemmli buffer with DTT and boiled for 5 min, samples were loaded on 4–20% Mini-PROTEAN® TGX™ Precast Protein Gels (Biorad). Proteins were transferred on PVDF membranes using a Transblot Turbo System (Biorad), blocked in 5% non-fat dry milk/PBS, and incubated overnight at 4°C with primary antibodies. After washing, membranes were incubated for 1 hour with secondary antibodies (Abcam). Images were acquired using a Fujifilm LAS 3000 Imager. Densitometric analysis were performed with ImageJ software (Fiji, RRID:SCR_002285). Band signal intensity was normalized for the respective loading control values (actin or SDHA).

Primary antibodies: HIF-1 alpha Antibody (Novus Cat# NB100-479SS, RRID:AB_790147), HIF2 alpha antibody (Abcam Cat# ab199, RRID:AB_302739), SDHA antibody (Abcam Cat# ab14715, RRID:AB_301433), 6X His tag® antibody (Abcam Cat# ab9108, RRID:AB_307016), beta Actin antibody (Abcam Cat# ab75186, RRID:AB_1280759). Secondary antibodies: Goat anti-Rabbit IgG (H+L) Secondary Antibody, HRP (Thermo Fisher Scientific Cat# 31460, RRID:AB_228341), Goat anti-Mouse IgG (H+L) Secondary Antibody, HRP (Thermo Fisher Scientific Cat# 31430, RRID:AB_228307).

### Colony and sphere formation assays

Mechanical dissociation with Accutase (ThermoFisher) was used to prepare single cell suspensions. Cells were counted using a hemocytometer. For colony formation assay, 350 cells were seeded in 5 cm dishes coated with polyornithine (Sigma) and laminin (Biolamina). U3082MG, U3084MG and U3065MG cells were cultured for 14 days while PIGPCs for 8 days, under the indicated conditions, then washed in PBS and fixed using 4% PFA. Cells were stained using 0,01% crystal violet/H2O. Wells were washed gently in water, then air-dried for 24 hours. Images were acquired with a Fujifilm LAS 3000 Imager.

Sphere formation assay was performed with the hanging-drop method. 10 cells in 35 µl drops were seeded on the lid of a 48 well plate and grown under the indicated conditions for 2 weeks. For secondary sphere assay, primary spheres were pooled, pelleted, dissociated with Accutase and reseeded at the indicated conditions. Wells with spheres were manually counted and images were acquired with a Zeiss AX10 inverted microscope.

### Real-Time Quantitative PCR

The RNeasy Mini Kit was used with Qiashredder (QIAGEN) according to the manufacturer’s instructions for RNA isolation, and cDNA was synthesized using random primers and Multiscribe reverse transcriptase (Applied Biosystems). A QuantStudio 7 real-time PCR system (Applied Biosystems) with SYBR Green Master Mix (Applied Biosystems) was used for amplification. Gene expression levels were normalized to the expression of three housekeeping genes (UBC, SDHA, and YWHAZ) using the comparative ΔΔCT method.

The following primers were used: *NANOG* (GCTGGTTGCCTCATGTTATTATGC; CCACATCGGCCTGTGTATATC), *UBC* (ATTTGGGTCGCGGTTCTT; TGCCTTGACATTCTCGATGGT), *SDHA* (TGGGAACAAGAGGGCATCTG; CCACCACTGCATCAAATTCATG), *YWHAZ* (ACTTTTGGTACATTGTGGCTTCAA; CCGCCAGGACAAACCAGTAT)

### Transfections and luciferase reporter assay

DLK-S expression plasmid was kindly provided by prof. Anne Ferguson-Smith (Ferrón et al., 2011), cloned into RCAS vector by classic restriction enzyme technique and transfected into DF-1 cells. For luciferase reporter assay, cells were co-transfected with HRE-luc (Addgene) (Emerling, Weinberg, Liu, Mak, & Chandel, 2008) or 8xCSL-luc (gift from Håkan Axelson) and pCMV-renilla (Promega) and analyzed using the Dual-Luciferase Reporter Assay System (Promega) on a Synergy 2 platereader (BioTek). Xtreme gene 9 (Roche) reagent was used according to manufacturer’s recommendations for transient transfections.

### Statistical Analyses

For DLK1 expression and patients survival, data from 620 patients from The Cancer Genome Atlas (TCGA, RRID:SCR_003193, https://portal.gdc.cancer.gov/) (Ceccarelli et al., 2016), were analyzed using GlioVis ((http://gliovis.bioinfo.cnio.es/) (Bowman, Wang, Carro, Verhaak, & Squatrito, 2017)) (DLK1-LOW n=333, events=82, median=87.5; DLK1-HIGH n=334, events=157, median=34.9).

For proliferation experiments, the EC50 and span were estimated with a nonlinear regression curve, using a log. agonist vs. normalized response (variable slope) equation fitted in Graphpad Prism 5 (GraphPad Prism, RRID:SCR_002798). For immunofluorescence experiments, colocalization was measure by Pearson’s R coefficient in 3 independent experiments with ImageJ Coloc2 plugin. For Ki67 quantification, at least 3 fields were analyzed for each tumor, for a total of 101717 nuclei analyzed, with n=45466 for PDGFB tumors and n=56251 for DLK-S tumors.

After normal distribution and variance similarity evaluation, two-sided unpaired t-test (eventual Welch’s correction for groups with different variances), Mann-Whitney for non-parametric data, one-way ANOVA with Bonferroni post-hoc test and two-way ANOVA (timelines only) tests were used to determine statistical significance, as indicated in respective figure legends. For survival evaluation, the Kaplan–Meier method was used to investigate variables and overall survival correlation, while a log-rank test was employed to compare survival curves. In all figures data are shown as mean±SEM, analyzed using GraphPad Prism 5 software and significance expressed as P values (* p < 0.05, ** p< 0.01, *** p < 0.001).

### Data availability

The data that support the findings of this study are available from the corresponding author upon reasonable request.

## RESULTS

### DLK1 is expressed and secreted by tumor-associated astrocytes in the glioma microenvironment

To test whether tumor-associated astrocytes could be a source of DLK1 in the glioma tumor microenvironment, we generated PDGFB/shp53-induced murine gliomas using the RCAS/tv-a system as previously described (Grassi et al., 2020), then co-stained tumors for the astrocyte marker GFAP and DLK1. While the bulk of the tumor cells appeared negative for DLK1 expression, DLK1 signal was detected in areas of GFAP staining both in perinecrotic and perivascular tumor areas, and both in GFAP positive and negative cells, suggesting DLK1 expression in astrocytes and tumor cells in these areas (Fig. 1A-B).

The use of an independent antibody directed against the N-terminal, soluble domain of DLK1 also showed specific signal in the perinecrotic tumor areas (Supp. Fig. 1) and co-staining of GFAP and DLK1 (Fig. 1C-D).

DLK1 gene expression was previously reported to be upregulated in isolated GFAP+ tumor-associated astrocytes of high-grade glioma compared to those of lower-grade tumors (Katz et al., 2012), further confirming DLK1 expression in astrocytes in this model system. We speculated that this DLK1 induction could be mediated by microenvironmental factors. As hypoxia is one major microenvironmental reality of high-grade glioma compared to low-grade glioma (Jawhari et al., 2016; Jensen, 2009), we cultured human fetal astrocytes under normoxic or hypoxic conditions for up to 10 days. We also subjected astrocytes to a single dose of irradiation to mimic another physiological response to a therapeutic intervention relevant to high-grade glioma. Both astrocytes subjected to growth in hypoxia and those subjected to irradiation displayed increased DLK1 levels secreted into the culture media, as measured by an ELISA assay (Fig. 2A). The higher levels of DLK1 in the media were sustained for the entire 10 day period in the case of astrocytes cultured in hypoxia, whereas irradiated astrocytes displayed a peak secretion at 2 days post-treatment with levels returning to baseline after 9 days (Fig. 2A). Together, these data support that astrocytes may secrete DLK1 into the tumor microenvironment in glioma.

**Figure 2.**
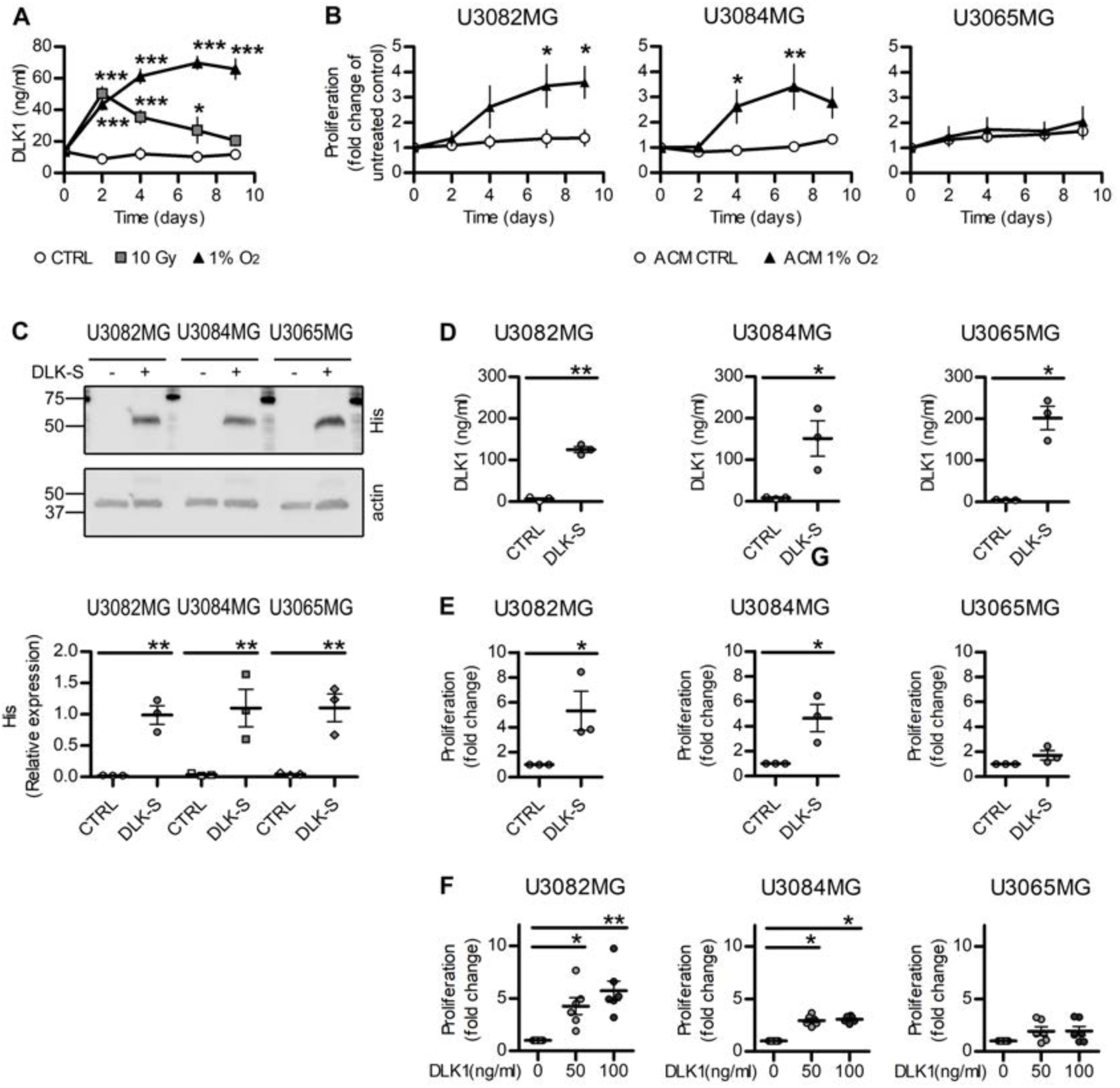
A: ELISA assay data showing DLK1 secretion in human fetal astrocytes exposed to one-time 10Gy irradiation or 1% O_2_ for up to 9 days. B: Media transfer experiments showing the effects of normoxic and hypoxic Astrocyte-Conditioned media (ACM) on human glioma cell lines. C: Representative images and densitometric analysis of western blots showing His-tagged soluble DLK1 expression in U3082MG, U3084 and U3065 cells transient transfection. D: ELISA assay data showing DLK1 secretion in human glioblastoma cells transfected with soluble DLK1. E: Media transfer experiments showing the effects of media from glioblastoma cells overexpressing soluble DLK1 on human glioma cell lines themselves. F: Proliferation assays of glioma cells treated with recombinant DLK1 for 72 hours. Statistical analysis: A, B n=4 C, D, E n=3, F=6. Statistical significance was determined by two-way ANOVA (A, B) and t-test (C-F), in E and F Welch’s correction for unequal variances was applied. In the whole figure significance is represented as * p<0.05, **p<0.01 and *** p<0.001 vs. untreated controls.

### Soluble DLK1 promotes glioma cell proliferation, survival and self-renewal

We next examined what effect soluble DLK1 may have on glioma cells.

We first performed a media transfer experiment by treating human glioblastoma cell lines maintained in serum-free, stem cell-promoting conditions with media from astrocytes cultured in normoxic (ACM CTRL) and hypoxic (ACM 1% O_2_) conditions for up to 9 days. In 2 out of 3 cell lines, media from hypoxic astrocytes induced a significant increase in the proliferation rate (Fig. 2B). Since astrocyte conditioned media contains many other different factors that may influence cancer cell growth, we then directly investigated the effects of soluble DLK1 with two different approaches. First, we transiently transfected human glioblastoma cell lines with a plasmid containing the N-terminal soluble part of DLK1 (DLK-S, His tagged). All the three transfected cell lines overexpressed and secreted similar levels of DLK-S, as verified by western blot and ELISA experiments (Fig. 2C-D). A media transfer experiment showed that 2 out of 3 cell lines significantly increased their proliferation (Fig. 2E) when grown in DLK-S conditioned media. We then treated the human glioblastoma cell lines with a recombinant protein corresponding to the DLK1 secreted part, and once again, the two DLK-responding cell lines showed significant increase in their proliferation (Fig. 2F). Taken together, these data demonstrate that soluble DLK1 is able to induce glioma cell proliferation, irrespectively of its origin.

As the use of the recombinant protein allows for better control of DLK1 concentrations, we then moved forward with this approach. We first generated a dose-response curve in all 3 human glioblastoma cell lines and in PDGFB-induced glioma primary cultures (PIGPCs) derived from the glioma mouse model. All the cell lines, with the exception of the non-responding U3065MG, showed a dose dependent increase in cell proliferation, with a plateau obtained at 200 ng/ml DLK1 and EC50s in between 25 and 35 ng/ml (Fig 3A).

**Figure 3.**
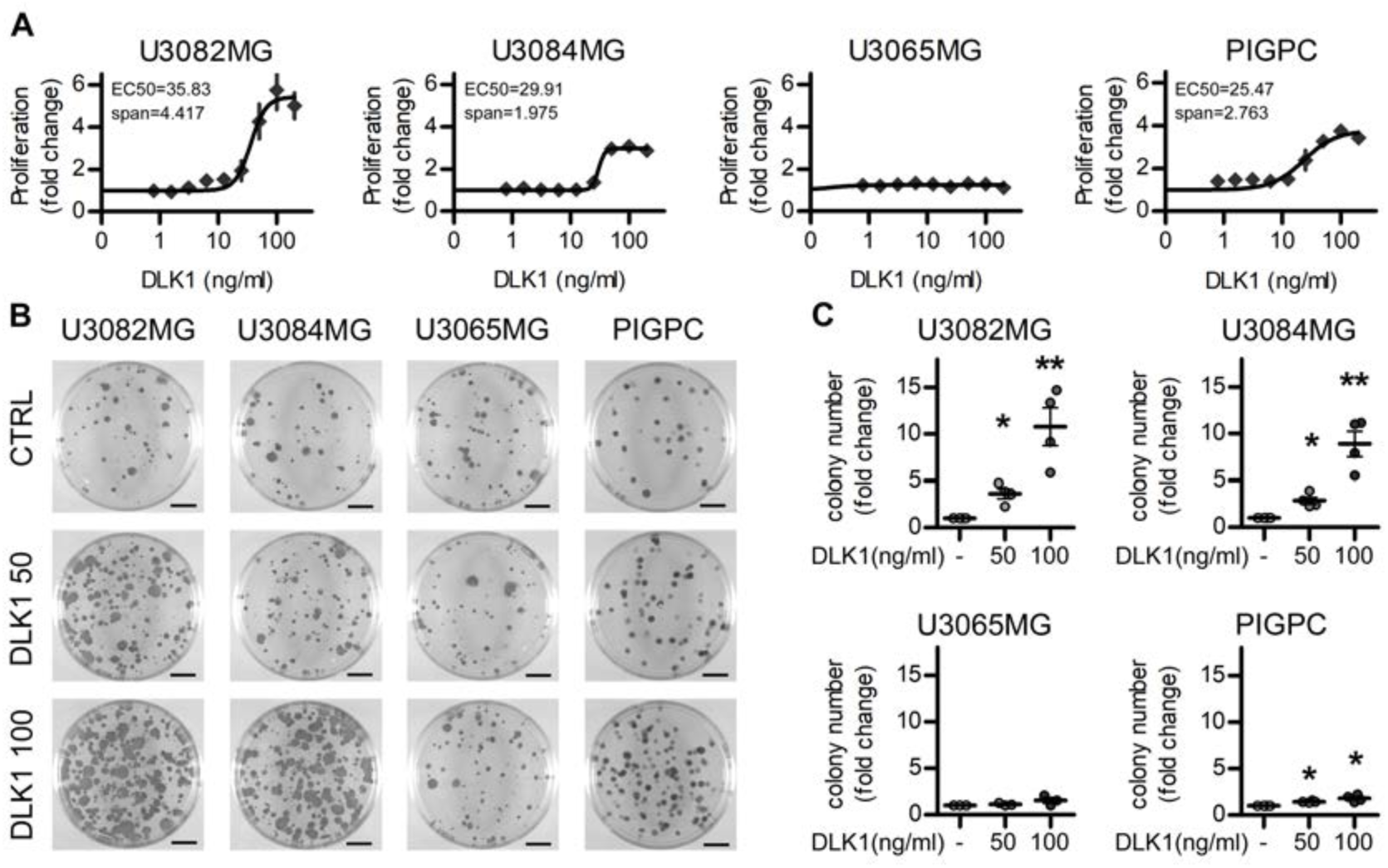
A: Proliferation curves of U3082MG, U3084MG, U3065MG and PIGPC cells grown at increasing concentrations of recombinant DLK1 for 72 hours. Data are expressed as fold change of untreated control. B, C: Representative images and quantification of colony forming ability of U3082MG, U3084MG, U3065MG and PIGPC cells grown at increasing concentrations of recombinant DLK1. Data are expressed as fold change of untreated control. Statistical analysis: independent experimental replicates are as follow, A n=6, except U3065MG where n=4, C n=4, Statistical significance was determined by one-way ANOVA, followed by Bonferroni post hoc test. In the whole figure significance is represented as * p<0.05 and ** p<0.01 vs. untreated controls.

In line with these findings, all cell lines that responded to DLK1 in the proliferation assay also increased their colony formation ability in a dose-dependent manner when exposed to sub-maximal soluble DLK1 concentrations similar to those obtained in hypoxic astrocytes (Fig. 3B-C).

Furthermore, soluble DLK1 strongly enhanced the self-renewal ability of responsive glioma cell lines, as measured by the serial sphere-formation assay (Fig. 4A-B), and induced a significant increase in the stemness markers *OCT4, NANOG* and *SOX2* (Fig. 4C). Since DLK1 has been reported to influence the Notch pathway (Baladrón et al., 2005; Huang et al., 2019), we tested if these effects were Notch-dependent. Luciferase experiments performed at different time points revealed no significant alterations in Notch activity in any tested cell lines (Fig. 4D).

**Figure 4.**
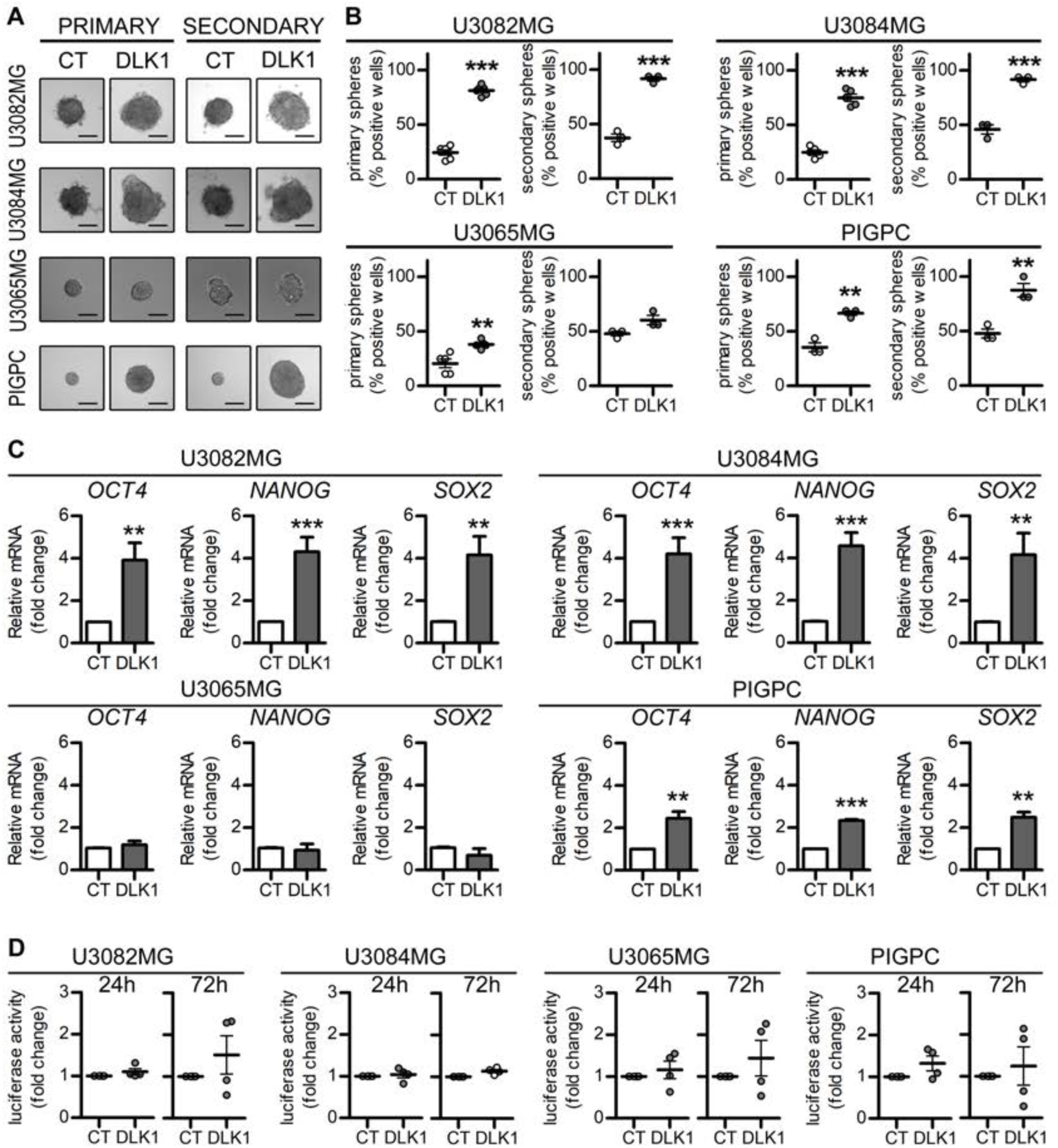
A, B: Representative images and quantification of primary and secondary sphere forming assays in U3082MG, U3084MG, U3065MG and PIGPC cells grown at 0 or 50 ng/ml recombinant DLK1. C: qPCR data for relative mRNA expression of *OCT4, NANOG* and *SOX2* in U3082MG, U3084MG, U3065MG and PIGPC cells grown at 0 or 50 ng/ml recombinant DLK1 for 72 hours. Data are expressed as fold change of untreated control. D: 8xCSL-luciferase experiments showing Notch activity in cell lines untreated or treated with 50 ng/ml recombinant DLK1 for 24 and 72 hours. Results are expressed as fold change of respective controls. Statistical analysis: independent experimental replicates are as follow, B n=5 for U3082MG, U3084MG and U3065MG primary spheres and n=3 PIGPC cells, C n=8 except PIGPC where n=3, D n=4. Statistical significance was determined by t-test, in C and D Welch’s correction for unequal variances was applied. In the whole figure significance is represented as * p<0.05, ** p<0.01 and *** p<0.001 vs. untreated controls.

### DLK1-effects are mediated in part by HIF-2alpha

Because DLK1 secretion was increased by astrocytes under hypoxic conditions, and because of known previous links between DLK1 expression and function to hypoxia (Kim et al., 2009), we asked whether soluble DLK1 could influence the hypoxic response of glioma cells. We cultured U3082MG and U3084MG human glioblastoma cells for 24 or 72 h in 1% O_2_, stimulated or not with soluble DLK1. While there was no difference in HIF-1alpha stabilization with DLK1 treatment, Western blots showed significantly increased HIF-2alpha protein levels at 72 h in cells cultured with soluble DLK1 (Fig. 5A-B). This increased HIF-2alpha expression was reflected in a stronger hypoxic response, as 2/3 human glioblastoma lines and PIGPCs displayed increased activation of hypoxia-responsive elements (HREs) at 72 h of culture in hypoxia with DLK1 stimulation, as measured in an HRE-luciferase assay (Fig. 5C). Moreover, analysis of PDGFB/shp53-induced murine gliomas revealed that both DLK1 and HIF-2alpha were strongly expressed and showed a significant co-localization only in the perivascular and perinecrotic niches (Fig 5D).

**Figure 5.**
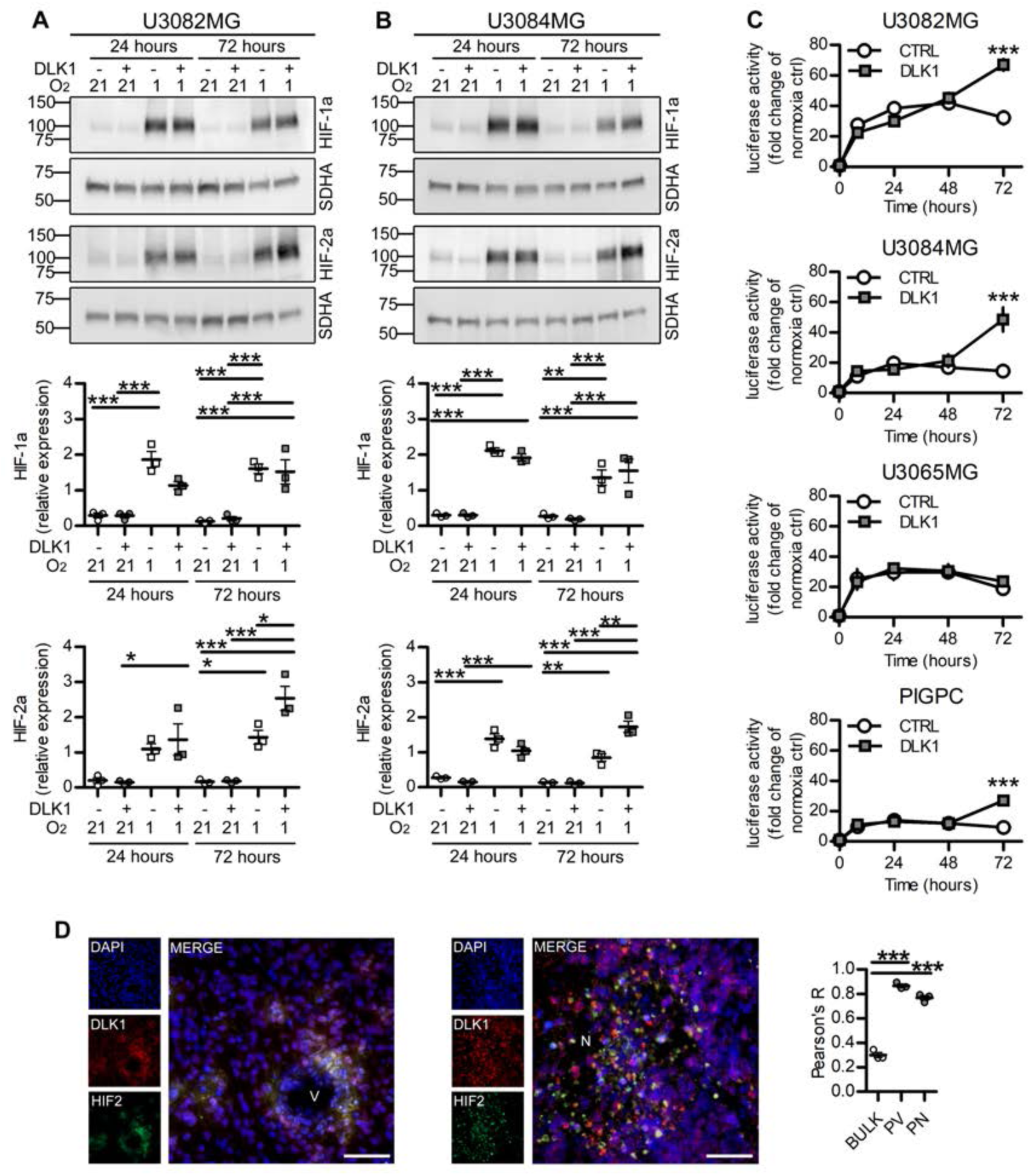
A: Representative images and densitometric analysis of western blots showing HIF-1alpha (HIF1-a) and HIF-2alpha (HIF2-a) expression in U3082MG cells after treatment with 50 ng/ml recombinant DLK1 or hypoxia exposure as indicated in the figure. B: : Representative images and densitometric analysis of western blots showing HIF-1alpha (HIF1-a) and HIF-2alpha (HIF2-a) expression in U3084MG cells after treatment with 50 ng/ml recombinant DLK1 or hypoxia exposure as indicated in the figure. C: HRE-luciferase time course experiments showing hypoxia response in cell lines untreated or treated with 50 ng/ml recombinant DLK1 and grown in 1% O_2_ for up to 72 hours. Results are expressed as fold changes of respective normoxic controls. D: Representative images and colocalization analysis of immunofluorescent staining showing DLK1 and HIF2-alpha (HIF2) expression in perivascular and hypoxic niches. V, vessel; N, necrosis. Scale bars represent 50 µm. Statistical analysis: independent experimental replicates are as follow, A, B n=3, C n=5 for U3082MG and n=4 for the other cell lines, D n=3. Statistical significance was determined by one-way ANOVA (A, B) followed by Bonferroni post hoc test, two-way ANOVA (C), one-way ANOVA of Pearson’s coefficients (D). In the whole figure significance is represented as * p<0.05, ** p<0.01 and *** p<0.001 vs. control or as indicated by straight lines.

As HIF-2alpha is a known driver of stem cell characteristics in glioma and other tumor forms (Das et al., 2019; Johansson et al., 2017; Pietras et al., 2008; Pietras, Johnsson, & Påhlman, 2010; Yan et al., 2018), we next tested whether effects of DLK1 on glioma cell behavior were mediated by HIF-2alpha. The treatment with the specific HIF-2alpha inhibitor PT2385 at concentrations with significant effects on HIF-2alpha protein (Persson et al., 2020; Wallace et al., 2016) (Supp. Fig 2) was able to revert the DLK1-induced increase in hypoxia response (Fig. 6A). Similarly, while stimulation of glioma cells with soluble DLK1 boosted the increase in the expression of the stem cell marker genes *NANOG, OCT4* and *SOX2* in hypoxic cells, addition of PT2385 blocked this specific DLK1 effect in all cell lines tested (Fig. 6B). Moreover, PT2385 decreased the colony formation ability of glioma cells exposed to soluble DLK1 in hypoxia (Fig. 6C). Notably however, PT2385 treatment did not significantly affect DLK1-induced gene expression in normoxia. Together, these data suggest that DLK1 promotes the glioma stem cell character in part via HIF-2alpha stabilization.

**Figure 6.**
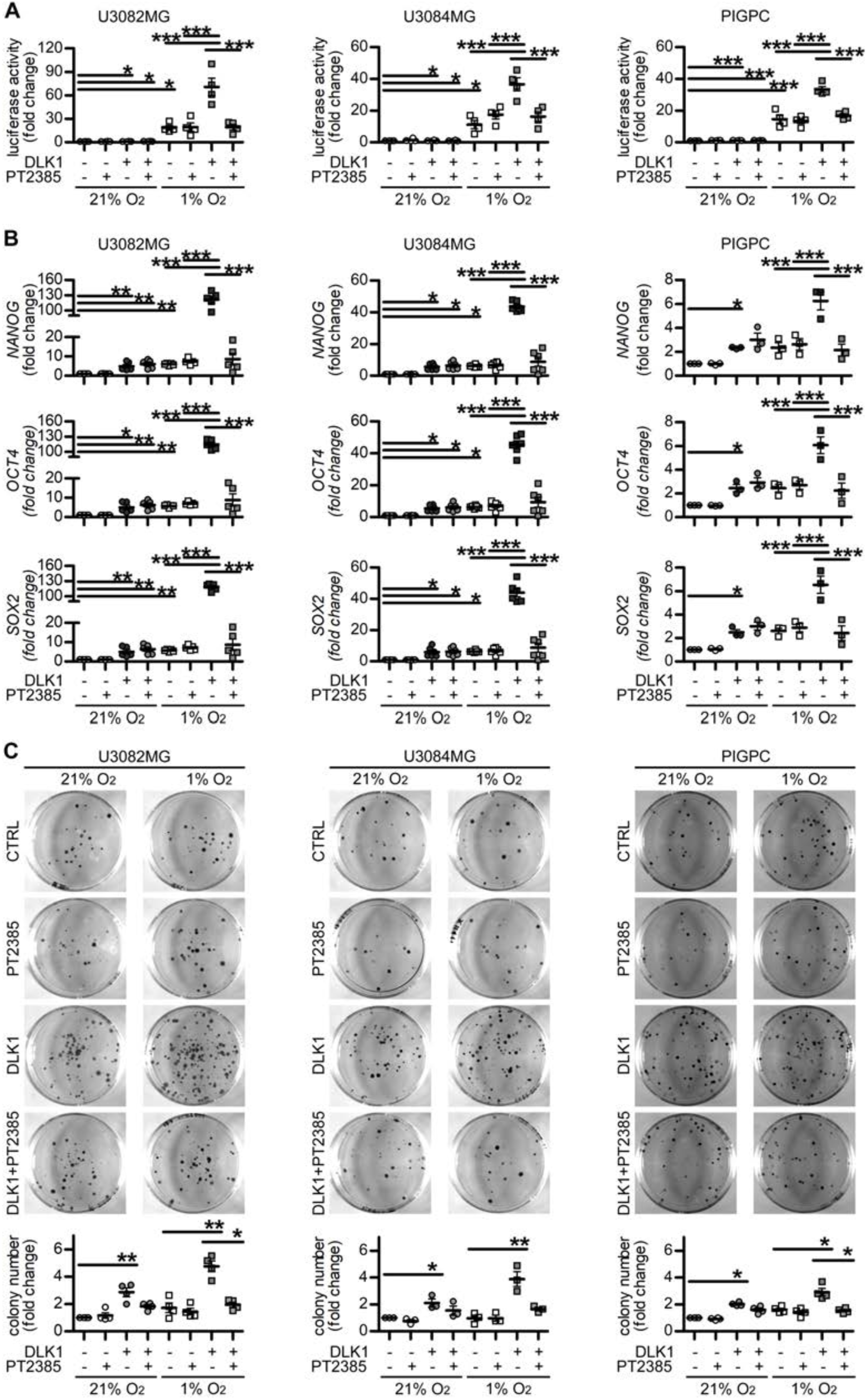
A: HRE-luciferase experiments showing hypoxia response in cell lines untreated or treated with 50 ng/ml recombinant DLK1, 10 µM PT2385 and grown in 21% or 1% O_2_ for 72 hours. Results are expressed as fold changes of respective normoxic controls. B: qPCR data for relative mRNA expression of *NANOG, OCT4* and *SOX2* in U3082MG, U3084MG and PIGPC cells pre-treated with 50 ng/ml recombinant DLK1, 10 µM PT2385 and exposed to hypoxia for 72 hours as indicated in the figure. Data are expressed as fold change of untreated normoxic control. C: Representative images and quantification of colony forming ability of U3082MG, U3084MG and PIGPC cells pre-treated with 50 ng/ml recombinant DLK1, 10 µM PT2385 and exposed to hypoxia as indicated in the figure. Data are expressed as fold change of untreated normoxic control. Statistical analysis: independent experimental replicates are as follow, A n=4, B n=6 for U3082MG and U3084MG and n=3 for PIGPC, C n=4. All data are expressed as mean±SEM. Statistical significance was determined by one-way ANOVA followed by Bonferroni post hoc test. In the whole figure significance is represented as * p<0.05, ** p<0.01 and *** p<0.001 as indicated by straight lines.

### DLK1 promotes aggressive glioma growth in vivo

To test the effects of soluble DLK1 on glioma growth in vivo, we generated a mouse model for the overexpression of soluble DLK1 together with PDGFB using the RCAS/tv-a system (Fig. 7A). Co-injection of RCAS-PDGFB with RCAS-DLK-S (soluble) resulted in more aggressive tumors as compared to RCAS-PDGFB with empty vector control, as measured by survival time following injections (Fig. 7B). Evaluation of Ki67 expression revealed a significant increase in cell proliferation in murine DLK-S tumors as compared to controls (Fig. 7C, D), thus confirming the *in vitro* data (Fig. 2E-F, 3A). In agreement with our in vivo data, analysis of the human TCGA GBMLGG dataset (Ceccarelli et al., 2016) revealed that tumors expressing high levels of *DLK1* were significantly more aggressive than those with low levels of *DLK1* (Fig. 7E), presumably as a result of the higher *DLK1* levels reported in high-grade glioma.

**Figure 7.**
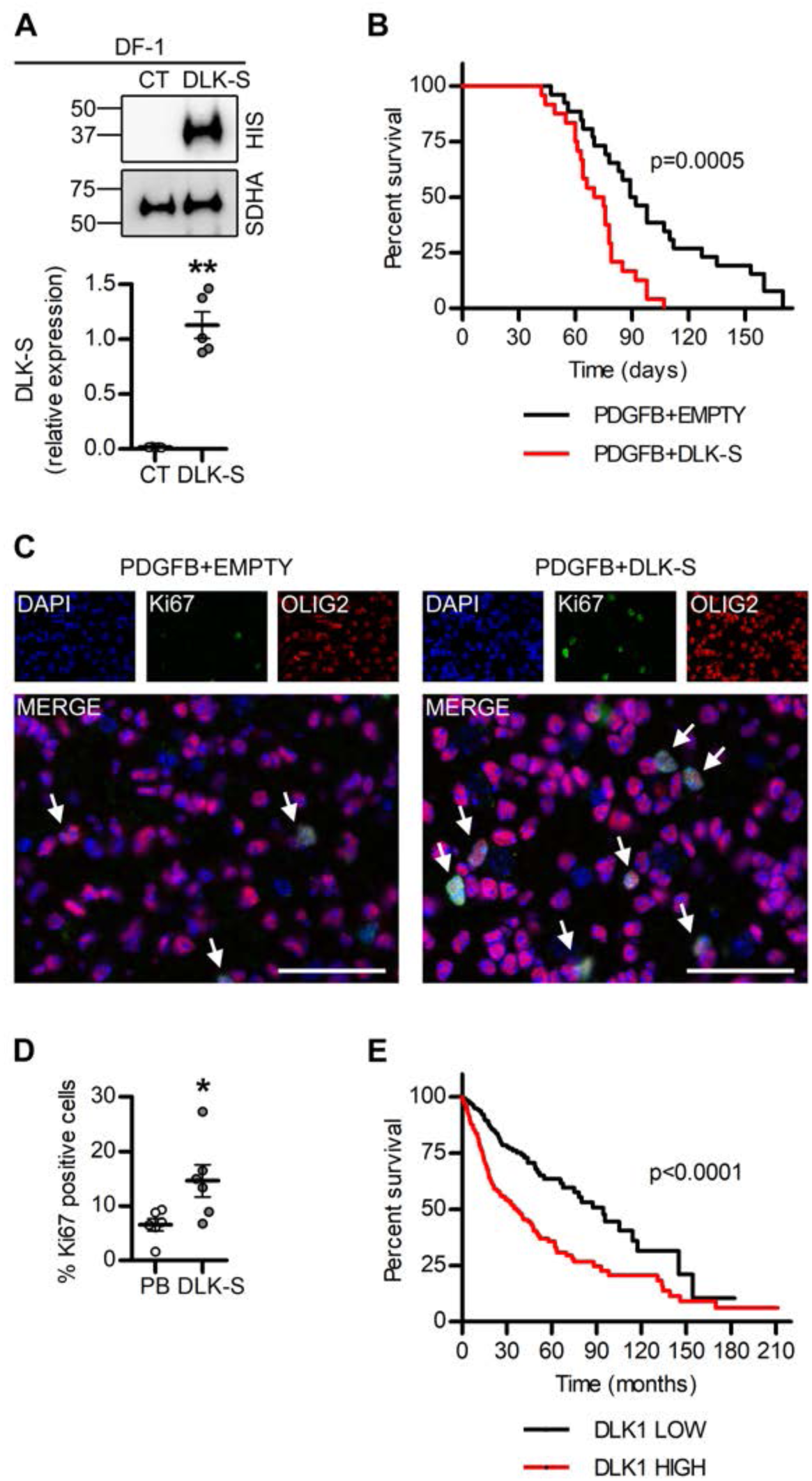
A: Representative images and densitometric analysis of western blots showing HIS-tagged DLK-S expression in DF1 cells transfected with empty or DLK-S RCAS vectors. B: Kaplan-Meier survival plot of PDGFB-induced tumors with (DLK-S, red line) or without (EMPTY, black line) DLK1 overexpression. C, D: Representative images and quantification of immunofluorescent stainings showing Ki67 positive cells expressed as percentage of tumor cells identified by OLIG2 staining. Scalebars represents 50 µm. E: Kaplan-Meier survival plot of DLK1-HIGH (red line) and DLK1-LOW (black line) patients in TCGA GBMLGG dataset Statistical analysis: independent experimental replicates are as follow, A n=4, B n=21 for PDGFB and n=27 for DLK-S, C n=6, E n=333, events=82 for DLK1-low and n=334, events=157 for DLK1-high. Statistical significance was determined by t-test with Welch’s correction (A), Kaplan-Meier survival plot with log-rank Mantel-Cox test (B, C) and Mann-Whitney (D). In the whole figure significance is represented as * p<0.05, ** p<0.01 vs. control.

## DISCUSSION

An increasing focus on cancer stemness has revealed parallels between normal neural stem cell regulation and cancer stem cell characteristics in brain tumors (Dirks, 2010; Lathia, Mack, Mulkearns-Hubert, Valentim, & Rich, 2015). Control over tumor cell phenotypes by specific, local microenvironments within a tumor, for example, is reminiscent of the way that normal tissue stem cells reside within and rely on their niche to maintain the stem cell character (Dirkse et al., 2019; Hambardzumyan & Bergers, 2015). It is likely that some of the same mechanisms involved in neural stem cell maintenance in the vascular niche of the SVZ may also be involved in maintaining stemness of brain tumor cells located in a perivascular niche. We describe one such example here: soluble DLK1 secreted from astrocytes appears to be involved in stemness maintenance both of normal neural stem cells and glioma cells, as shown here. An association between DLK1 expression and aggressive tumor growth in glioblastoma has previously been established (Kim et al., 2009). By generating a mouse model for testing effects of soluble DLK1 overexpression specifically in the context of glioblastoma, we show that the previously reported association between DLK1 expression and tumor grade in glioma (Grassi et al., 2020, Yin et al., 2006) may at least in part be caused by DLK1 itself, as soluble DLK1-overexpressing tumors had a higher proliferation rate and significantly decreased mice survival as compared to controls. It is important to note that DLK1 expression is not limited to tumor-associated astrocytes, and soluble DLK1 may be derived both from other stromal cell types and tumor cells themselves. Our experiments indicate that soluble DLK1 affects tumor cell proliferation similarly regardless of whether it was produced by astrocytes or tumor cells.

As with astrocyte-derived DLK1 in regulation of normal neural stem cells, the exact mechanism(s) by which soluble DLK1 signals to glioma cells remains to be investigated. We show here that soluble DLK1 can contribute to a stronger and more prolonged response to hypoxia, as mediated by increased HIF-2alpha stabilization in DLK1 treated cells. This effect on HIF-2alpha stabilization indeed seemed important for the tumor-promoting effects of DLK1 signaling as inhibition of HIF-2alpha transcriptional activity by use of the specific HIF-2alpha inhibitor PT2385 abolished all effects of DLK1 on stem cell marker gene expression and colony formation under hypoxic conditions. Interestingly, DLK1 expression itself has been shown to be regulated by hypoxia in other cell systems (Begum et al., 2012; Kim et al., 2009), suggesting that there may be a DLK1-HIF feedback loop in hypoxic tumor cells. In the present investigation, effects of DLK1 treatment were enhanced by hypoxic culture conditions. Importantly, however, HIF-2alpha inhibition did not significantly affect DLK1-mediated stem cell marker expression under normoxic conditions, suggesting that there are other mediators downstream of DLK1 that can contribute to DLK1 signaling in glioma.

Taken together, our data support a role for soluble DLK1 as a tumor-promoting stem cell niche factor in glioma. Further research is warranted to investigate whether or not signaling by DLK1 can be therapeutically targeted, either via HIF-2alpha inhibition or by targeting upstream signaling.

## Acknowledgements

The authors thank Christina Möller for technical assistance and A Human Glioblastoma Cell Culture Resource (www.hgcc.se) (Lene Uhrbom, Bengt Westermark, Karin Forsberg Nilsson and Sven Nelander, Uppsala University, Sweden) for human GBM cultures. This work was supported by grants to AP from the Ragnar Söderberg Foundation, the Swedish Cancer Society, the Swedish Research Council, the Swedish Childhood Cancer Fund, Ollie & Elof Ericssons foundation, Jeanssons stiftelser, the Crafoord foundation, Gösta Miltons donationsfond, and Stiftelsen Cancera.

## REFERENCES

Baladrón, V., Ruiz-Hidalgo, M. J., Nueda, M. L., Díaz-Guerra, M. J. M., García-Ramírez, J. J., Bonvini, E., … Laborda, J. (2005). dlk acts as a negative regulator of Notch1 activation through interactions with specific EGF-like repeats. Experimental Cell Research, 303(2), 343–59. https://doi.org/10.1016/j.yexcr.2004.10.001

Begum, A., Kim, Y., Lin, Q., & Yun, Z. (2012). DLK1, delta-like 1 homolog (Drosophila), regulates tumor cell differentiation in vivo. Cancer Letters, 318(1), 26–33. https://doi.org/10.1016/j.canlet.2011.11.032

Bernstock, J. D., Mooney, J. H., Ilyas, A., Chagoya, G., Estevez-Ordonez, D., Ibrahim, A., & Nakano, I. (2019). Molecular and cellular intratumoral heterogeneity in primary glioblastoma: clinical and translational implications. Journal of Neurosurgery, 1–9. https://doi.org/10.3171/2019.5.JNS19364

Bleau, A.-M., Hambardzumyan, D., Ozawa, T., Fomchenko, E. I., Huse, J. T., Brennan, C. W., & Holland, E. C. (2009). PTEN/PI3K/Akt Pathway Regulates the Side Population Phenotype and ABCG2 Activity in Glioma Tumor Stem-like Cells. Cell Stem Cell, 4(3), 226–235. https://doi.org/10.1016/j.stem.2009.01.007

Bowman, R. L., Wang, Q., Carro, A., Verhaak, R. G. W., & Squatrito, M. (2017). GlioVis data portal for visualization and analysis of brain tumor expression datasets. Neuro-Oncology, 19(1), 139–141. https://doi.org/10.1093/neuonc/now247

Ceccarelli, M., Barthel, F. P., Malta, T. M., Sabedot, T. S., Salama, S. R., Murray, B. A., … Verhaak, R. G. W. (2016). Molecular Profiling Reveals Biologically Discrete Subsets and Pathways of Progression in Diffuse Glioma. Cell, 164(3), 550–563. https://doi.org/10.1016/j.cell.2015.12.028

Ceder, J. A., Jansson, L., Helczynski, L., & Abrahamsson, P.-A. (2008). Delta-like 1 (Dlk-1), a novel marker of prostate basal and candidate epithelial stem cells, is downregulated by notch signalling in intermediate/transit amplifying cells of the human prostate. European Urology, 54(6), 1344–53. https://doi.org/10.1016/j.eururo.2008.03.006

Das, B., Pal, B., Bhuyan, R., Li, H., Sarma, A., Gayan, S., … Felsher, D. W. (2019). MYC Regulates the HIF2α Stemness Pathway via Nanog and Sox2 to Maintain Self-Renewal in Cancer Stem Cells versus Non-Stem Cancer Cells. Cancer Research, 79(16), 4015–4025. https://doi.org/10.1158/0008-5472.CAN-18-2847

Dirks, P. B. (2010, October 1). Brain tumor stem cells: The cancer stem cell hypothesis writ large. Molecular Oncology. John Wiley and Sons Ltd. https://doi.org/10.1016/j.molonc.2010.08.001

Dirkse, A., Golebiewska, A., Buder, T., Nazarov, P. V., Muller, A., Poovathingal, S., … Niclou, S. P. (2019). Stem cell-associated heterogeneity in Glioblastoma results from intrinsic tumor plasticity shaped by the microenvironment. Nature Communications, 10(1), 1787. https://doi.org/10.1038/s41467-019-09853-z

Emerling, B. M., Weinberg, F., Liu, J.-L., Mak, T. W., & Chandel, N. S. (2008). PTEN regulates p300-dependent hypoxia-inducible factor 1 transcriptional activity through Forkhead transcription factor 3a (FOXO3a). Proceedings of the National Academy of Sciences of the United States of America, 105(7), 2622–7. https://doi.org/10.1073/pnas.0706790105

Falix, F. A., Aronson, D. C., Lamers, W. H., & Gaemers, I. C. (2012). Possible roles of DLK1 in the Notch pathway during development and disease. Biochimica et Biophysica Acta, 1822(6), 988–95. https://doi.org/10.1016/j.bbadis.2012.02.003

Ferrón, S. R., Charalambous, M., Radford, E., McEwen, K., Wildner, H., Hind, E., … Ferguson-Smith, A. C. (2011). Postnatal loss of Dlk1 imprinting in stem cells and niche astrocytes regulates neurogenesis. Nature, 475(7356), 381–385. https://doi.org/10.1038/nature10229

Grassi, E. S., Pantazopoulou, V., & Pietras, A. (2020). Hypoxia-induced release, nuclear translocation, and signaling activity of a DLK1 intracellular fragment in glioma. Oncogene, 1–17. https://doi.org/10.1038/s41388-020-1273-9

Hambardzumyan, D., & Bergers, G. (2015). Glioblastoma: Defining Tumor Niches. Trends in Cancer, 1(4), 252–265. https://doi.org/10.1016/j.trecan.2015.10.009

Holland, E. C., Hively, W. P., DePinho, R. A., & Varmus, H. E. (1998). A constitutively active epidermal growth factor receptor cooperates with disruption of G1 cell-cycle arrest pathways to induce glioma-like lesions in mice. Genes & Development, 12(23), 3675–3685. https://doi.org/10.1101/gad.12.23.3675

Huang, C.-C., Cheng, S.-H., Wu, C.-H., Li, W.-Y., Wang, J.-S., Kung, M.-L., … Tai, M.-H. (2019). Delta-like 1 homologue promotes tumorigenesis and epithelial-mesenchymal transition of ovarian high-grade serous carcinoma through activation of Notch signaling. Oncogene, 38(17), 3201–3215. https://doi.org/10.1038/s41388-018-0658-5

Huse, J. T., & Holland, E. C. (2010). Targeting brain cancer: advances in the molecular pathology of malignant glioma and medulloblastoma. Nature Reviews Cancer, 10(5), 319–331. https://doi.org/10.1038/nrc2818

Jawhari, S., Ratinaud, M.-H., & Verdier, M. (2016). Glioblastoma, hypoxia and autophagy: a survival-prone “ménage-à-trois”. Cell Death & Disease, 7(10), e2434. https://doi.org/10.1038/cddis.2016.318

Jensen, R. L. (2009, May). Brain tumor hypoxia: Tumorigenesis, angiogenesis, imaging, pseudoprogression, and as a therapeutic target. Journal of Neuro-Oncology. Kluwer Academic Publishers. https://doi.org/10.1007/s11060-009-9827-2

Johansson, E., Grassi, E. S., Pantazopoulou, V., Tong, B., Lindgren, D., Berg, T. J., … Pietras, A. (2017). CD44 Interacts with HIF-2α to Modulate the Hypoxic Phenotype of Perinecrotic and Perivascular Glioma Cells. Cell Reports, 20(7). https://doi.org/10.1016/j.celrep.2017.07.049

Katz, A. M., Amankulor, N. M., Pitter, K., Helmy, K., Squatrito, M., & Holland, E. C. (2012). Astrocyte-specific expression patterns associated with the PDGF-induced glioma microenvironment. PLoS ONE, 7(2), e32453. https://doi.org/10.1371/journal.pone.0032453

Kim, Y., Lin, Q., Zelterman, D., & Yun, Z. (2009). Hypoxia-regulated delta-like 1 homologue enhances cancer cell stemness and tumorigenicity. Cancer Research, 69(24), 9271–80. https://doi.org/10.1158/0008-5472.CAN-09-1605

Lathia, J. D., Mack, S. C., Mulkearns-Hubert, E. E., Valentim, C. L. L., & Rich, J. N. (2015, June 15). Cancer stem cells in glioblastoma. Genes and Development. Cold Spring Harbor Laboratory Press. https://doi.org/10.1101/gad.261982.115

Li, L., Tan, J., Zhang, Y., Han, N., Di, X., Xiao, T., … Liu, Y. (2014). DLK1 promotes lung cancer cell invasion through upregulation of MMP9 expression depending on Notch signaling. PloS One, 9(3), e91509. https://doi.org/10.1371/journal.pone.0091509

Majmundar, A. J., Wong, W. J., & Simon, M. C. (2010). Hypoxia-inducible factors and the response to hypoxic stress. Molecular Cell, 40(2), 294–309. https://doi.org/10.1016/j.molcel.2010.09.022

Mega, A., Hartmark Nilsen, M., Leiss, L. W., Tobin, N. P., Miletic, H., Sleire, L., … Östman, A. (2020). Astrocytes enhance glioblastoma growth. GLIA, 68(2), 316–327. https://doi.org/10.1002/glia.23718

Persson, C. U., von Stedingk, K., Fredlund, E., Bexell, D., Påhlman, S., Wigerup, C., & Mohlin, S. (2020). ARNT-dependent HIF-2 transcriptional activity is not sufficient to regulate downstream target genes in neuroblastoma. Experimental Cell Research, 388(2), 111845. https://doi.org/10.1016/j.yexcr.2020.111845

Pietras, A., Gisselsson, D., Ora, I., Noguera, R., Beckman, S., Navarro, S., & Påhlman, S. (2008). High levels of HIF-2alpha highlight an immature neural crest-like neuroblastoma cell cohort located in a perivascular niche. The Journal of Pathology, 214(4), 482–8. https://doi.org/10.1002/path.2304

Pietras, A., Johnsson, A. S., & Påhlman, S. (2010). The HIF-2α-driven pseudo-hypoxic phenotype in tumor aggressiveness, differentiation, and vascularization. Current Topics in Microbiology and Immunology, 345, 1–20. https://doi.org/10.1007/82_2010_72

Ryskalin, L., Gaglione, A., Limanaqi, F., Biagioni, F., Familiari, P., Frati, A., … Fornai, F. (2019, August 1). The autophagy status of cancer stem cells in gliobastoma multiforme: From cancer promotion to therapeutic strategies. International Journal of Molecular Sciences. MDPI AG. https://doi.org/10.3390/ijms20153824

Sun Y, Vandenbriele C, Kauskot A, Verhamme P, Hoylaerts MF, Wright GJ. (2015). A human platelet receptor protein microarray identifies FcepsilonR1alpha as an activating PEAR1 ligand. Mol Cell Proteomics. Feb 23. pii: mcp.M114.046946.

Wallace, E. M., Rizzi, J. P., Han, G., Wehn, P. M., Cao, Z., Du, X., … Josey, J. A. (2016). A small-molecule antagonist of HIF2α is efficacious in preclinical models of renal cell carcinoma. Cancer Research, 76(18), 5491–5500. https://doi.org/10.1158/0008-5472.CAN-16-0473

Wang, Y., & Sul, H. S. (2006). Ectodomain shedding of preadipocyte factor 1 (Pref-1) by tumor necrosis factor alpha converting enzyme (TACE) and inhibition of adipocyte differentiation. Molecular and Cellular Biology, 26(14), 5421–35. https://doi.org/10.1128/MCB.02437-05

Xie, Y., Bergström, T., Jiang, Y., Johansson, P., Marinescu, V. D., Lindberg, N., … Uhrbom, L. (2015). The Human Glioblastoma Cell Culture Resource: Validated Cell Models Representing All Molecular Subtypes. EBioMedicine, 2(10), 1351–63. https://doi.org/10.1016/j.ebiom.2015.08.026

Yan, Y., Liu, F., Han, L., Zhao, L., Chen, J., Olopade, O. I., … Wei, M. (2018). HIF-2α promotes conversion to a stem cell phenotype and induces chemoresistance in breast cancer cells by activating Wnt and Notch pathways. Journal of Experimental & Clinical Cancer Research, 37(1), 256. https://doi.org/10.1186/s13046-018-0925-x

Yin, D., Xie, D., Sakajiri, S., Miller, C. W., Zhu, H., Popoviciu, M. L., … Koeffler, H. P. (2006). DLK1: increased expression in gliomas and associated with oncogenic activities. Oncogene, 25(13), 1852–61. https://doi.org/10.1038/sj.onc.1209219

