## Supplemental Data for "Niche-derived soluble DLK1 promotes glioma stemness and growth"

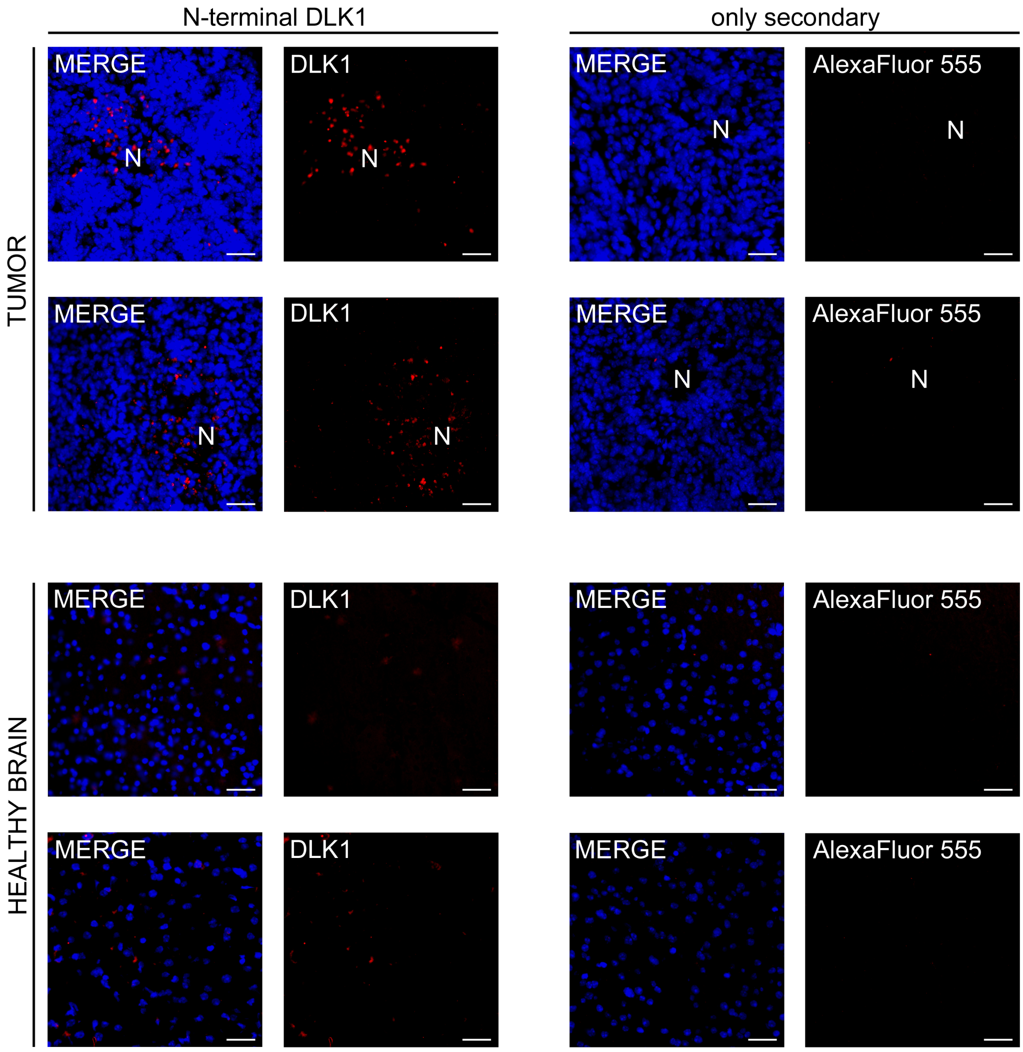


**Supp. figure 1**

Representative images of immunofluorescent stainings showing soluble DLK1 localization in shp53-induced murine gliomas and correspective healthy brain. Stainings with only secondaries only show no aspecific signal from N-terminal DLK1 antibody. Scalebars represents 25µm.

Statistical analysis: sections from 3 independent experiments were analyzed.

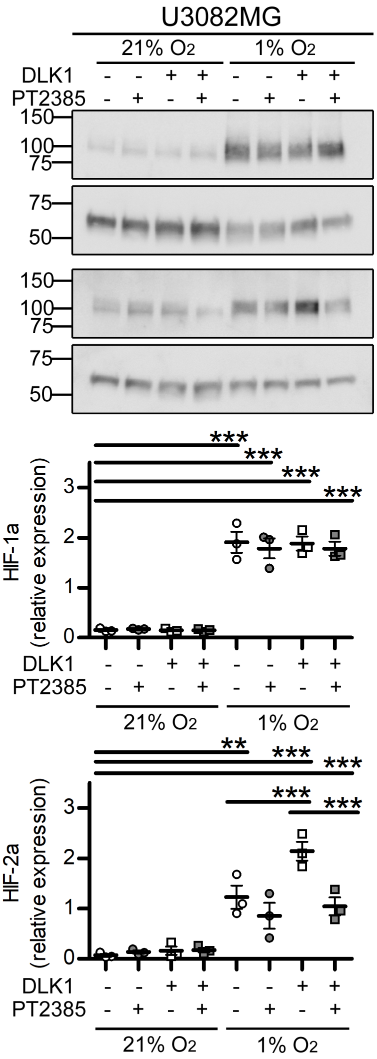


**Supp. figure 2**

Representative images and densitometric analysis of western blots showing HIF-1alpha (HIF1-a) and HIF-2alpha (HIF2-a) expression in U3082MG cells after treatment with 50 ng/ml recombinant DLK1, 10 µM PT2385 and hypoxia exposure as indicated in the figure.

Statistical analysis: n=3. All data are expressed as mean±SEM. Statistical significance was determined by one-way ANOVA followed by Bonferroni post hoc test. In the whole figure significance is represented as ** $p<0.01$ and *** $p<0.001$ as indicated by straight lines.
